# Enabling Spatially Resolved Gene Expression Analysis in a Multilayer Paper-Based *In Vitro* Tumor Model

**DOI:** 10.64898/2026.10.18.759958

**Authors:** Ankit Kumar, Bhushan J. Toley

## Abstract

Spatial heterogeneity is a defining feature of solid tumors, yet molecular analysis of three-dimensional (3D) *in vitro* tumor models is often performed as a bulk measurement, potentially obscuring position-dependent cellular responses. Here, we establish an analytical workflow for spatially resolved gene-expression analysis in a multilayer paper-based 3D tumor model by addressing the challenge of RNA recovery from cell-laden paper–Matrigel scaffolds. Using MDA-MB-231 breast cancer cells, multiple RNA extraction strategies were systematically evaluated across conventional culture, Matrigel, paper, and composite paper–Matrigel environments. Conventional silica-column extraction yielded only 3.39 ± 0.93 ng/µL RNA from the composite scaffold, whereas TRIzol–chloroform extraction increased recovery to 201.13 ± 38.32 ng/µL. The optimized workflow subsequently enabled independent RNA recovery from all eight layers of a multi-layer paper-based in vitro tumor, with concentrations ranging from 113.06 ± 5.00 to 245.26 ± 15.76 ng/µL and purity suitable for downstream qRT-PCR analysis. Layer-resolved profiling of untreated cultures revealed distinct spatial patterns in *HIF1A, HIF2A, VEGFA,* and *GLUT4* expression across the multilayer construct. To evaluate the workflow under pharmacological perturbation, cultures were exposed to 5 µM doxorubicin (DOX) for 48 h. Relative cell viability exhibited a non-monotonic spatial profile, whereas DOX-induced *HIF1A* and *VEGFA* expression generally became more pronounced toward the deeper layers when each treated layer was compared with its corresponding untreated control. Collectively, these results establish a workflow for converting individually addressable regions of a multilayer 3D tumor model into spatially resolved transcriptional measurements, enabling analysis of both baseline spatial heterogeneity and position-dependent molecular responses to pharmacological perturbation.

**Graphical Abstract:** 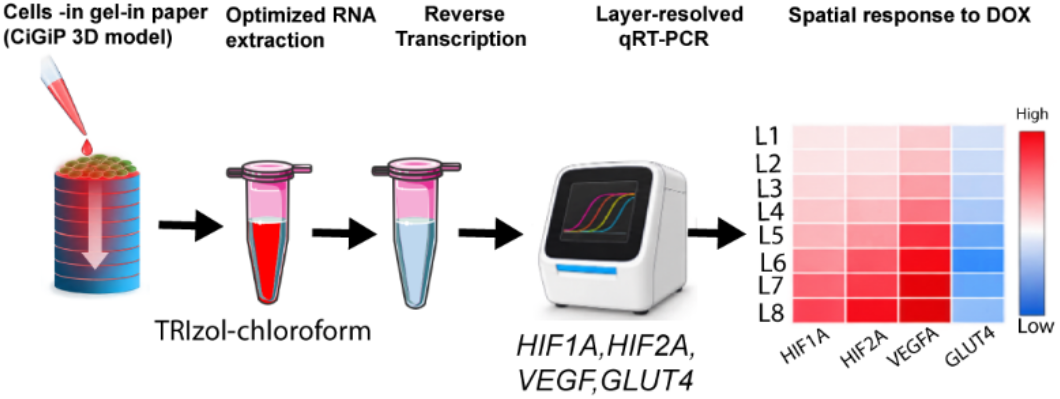

## Introduction

Three-dimensional (3D) cell culture systems have emerged as powerful in vitro models because they better reproduce the structural and biochemical complexity of native tissues than conventional two-dimensional (2D) monolayer cultures^1,2^. Unlike conventional 2D cultures, cells in 3D systems interact with neighbouring cells and the surrounding extracellular matrix (ECM). These interactions create gradients of oxygen, nutrients, metabolites, and signalling molecules that closely resemble the in vivo microenvironment^3–5^. As a result, 3D culture models have become valuable tools for studying cancer biology, drug screening, tissue engineering, and regenerative medicine^6–8^.

Among the various 3D culture approaches, paper-based scaffolds have attracted considerable attention because of their low cost, simplicity, biocompatibility, and ability to generate diffusion-limited microenvironments^9,10^. In these systems, porous cellulose paper serves as a structural scaffold, while hydrogels such as Matrigel provide an extracellular matrix that supports cell attachment, proliferation, and migration^11,12^. Stacking multiple cells-in-gel-in-paper (CiGiP) layers creates controllable oxygen and nutrient gradients that resemble those observed in avascular regions of solid tumors. Owing to their modular architecture, paper-based scaffolds also permit layer-by-layer separation, allowing spatial analysis of cellular behavior at different depths within the construct^13,14^.

One of the major advantages of multilayer paper-based culture systems is their potential to enable layer-specific molecular characterization, including gene expression analysis^15–17^. Cellular responses to oxygen and nutrient deprivation are regulated by hypoxia-inducible factors (*HIF-1α* and *HIF-2α*), angiogenic mediators such as vascular endothelial growth factor (*VEGF*), and glucose transporters including *GLUT4*^18–20^. Quantifying the expression of these genes across individual scaffold layers provides direct evidence of microenvironmental heterogeneity and cellular adaptation to diffusion-limited conditions. Although conventional 3D culture models, such as spheroids and hydrogels, are widely used, they offer limited access to cells within the construct, making it difficult to isolate cells from defined spatial regions and to recover high-quality RNA for downstream gene expression analysis. The layered architecture of paper-based culture systems offers an opportunity for spatially resolved molecular analysis because individual layers can be physically separated while retaining information about their position within the 3D construct. However, reliable recovery of RNA from cell-laden paper-based scaffolds remains comparatively underexplored. Although previous studies have demonstrated the potential of multilayer paper-based cultures for spatial analysis^21,22^, reports of layer-specific gene-expression measurements remain limited, and RNA extraction from these composite culture environments has not been systematically evaluated. Establishing a reliable RNA-recovery workflow is therefore important for extending physically addressable paper-based 3D cultures to layer-resolved transcriptional analysis.

In this study, we systematically evaluated RNA extraction strategies for a multilayer cells-in-gel-in-paper (CiGiP) 3D tumor model. The selected workflow was subsequently applied to investigate layer-dependent gene expression under baseline conditions and in response to doxorubicin treatment. Using MDA-MB-231 breast cancer cells, multiple RNA extraction strategies were systematically evaluated across conventional tissue culture, Matrigel, paper, and composite paper–Matrigel culture environments to identify the challenges associated with RNA recovery from increasingly complex 3D matrices. Based on comparative assessment of RNA yield, purity, and integrity, TRIzol–chloroform extraction method was selected as an optimized method for RNA recovery from individual, spatially defined layers of the multilayer construct. The resulting workflow was subsequently used to characterize baseline spatial transcriptional heterogeneity through layer-specific analysis of *HIF1A, HIF2A, VEGFA,* and *GLUT4*. To further demonstrate its analytical utility under pharmacological perturbation, the workflow was applied to doxorubicin-treated multilayer cultures, in which treatment-induced changes in *HIF1A* and *VEGFA* expression were quantified at each spatial position relative to the corresponding position in the untreated multilayer culture. Collectively, this study addresses a key sample-preparation barrier associated with molecular analysis of paper–Matrigel 3D cultures and establishes a workflow for converting physically addressable regions of a multilayer tumor model into spatially resolved transcriptional measurements. This capability expands the analytical utility of paper-based 3D culture platforms for investigating tumor microenvironmental heterogeneity and spatially dependent responses to anticancer treatment.

## Materials and Methods

### Chemicals and instruments

RNA extraction from MDA-MB-231 cells cultured in paper–Matrigel scaffolds was performed using either RNeasy Mini Kit (Qiagen, Hilden, Germany; Cat. No. 74104) or TRIzol™ Reagent (Thermo Fisher Scientific, Waltham, MA, USA; Cat. No. 15596018). The Qiagen kit contains RLT lysis buffer, RW1 wash buffer, RPE wash buffer, and RNeasy silica spin columns. Additional reagents used during RNA extraction and protocol optimization included chloroform (Sigma-Aldrich, St. Louis, MO, USA; Cat. No. C2432), isopropanol (Sigma-Aldrich, St. Louis, MO, USA; Cat. No. I9516), 70% ethanol (Sigma-Aldrich, St. Louis, MO, USA; Cat. No. E7023), β-Mercaptoethanol (Sigma-Aldrich, St. Louis, MO, USA), collagenase, trypsin–EDTA, and RNase-free water (Thermo Fisher Scientific, Waltham, MA, USA; Cat. No. AM9937).

RNA concentration and purity were determined using a NanoDrop™ 2000 spectrophotometer (Thermo Fisher Scientific, Waltham, MA, USA), while RNA integrity was evaluated by agarose gel electrophoresis. Complementary DNA (cDNA) was synthesized using the ProtoScript® II First Strand cDNA Synthesis Kit, New England Biolabs (NEB), Cat. No. E6560S. Quantitative real-time PCR was performed using the reagents and primers described in the corresponding methodology section. Mechanical homogenization was carried out using a Vortex-Genie 2 (Scientific Industries, Inc., Bohemia, NY, USA), and centrifugation was performed using an Eppendorf 5424R refrigerated microcentrifuge (Eppendorf AG, Hamburg, Germany).

### Cell culture

The human breast cancer cell line MDA-MB-231 was obtained as a kind gift from Prof. Ramray Bhat (Indian Institute of Science, Bengaluru, India). Cells were maintained in Dulbecco’s Modified Eagle Medium (DMEM) supplemented with 10% fetal bovine serum (FBS) and 1% antibiotic–antimycotic solution under standard culture conditions at 37 °C in a humidified atmosphere containing 5% CO₂. Cells were cultured in T25 culture flasks, and the culture medium was replaced every 2–3 days. Cells were subcultured at approximately 80–90% confluency using trypsin–EDTA and used for all experiments between appropriate passage numbers.

### Paper-based cell culture

#### Single-layer paper culture

Whatman Grade 1 filter paper was used as the scaffold for three-dimensional cell culture. Individual paper discs were sterilized prior to use by autoclaving followed by UV exposure. MDA-MB-231 cells were suspended at a density of 1 × 10⁷ cells mL⁻¹ and mixed with an equal volume of Matrigel while maintaining the mixture on ice to prevent premature gelation. Approximately 2 µL of cell suspension and 2 µL of Matrigel were carefully dispensed onto the center of each paper disc. The paper discs were incubated at 37 °C for 3 min to allow Matrigel polymerization. Subsequently, each paper disc was transferred to an individual well of a 12-well plate containing 250 µL of complete culture medium and incubated for 48 h before further processing.

#### Multilayer paper culture

For multilayer culture, individual paper discs containing CiGiP were stacked inside a previously developed 3D-printed press-fit culture device to generate a multilayer tissue construct^23^. Each scaffold consisted of eight paper layers (L1–L8), with each paper layer having a thickness of approximately 200 µm, resulting in a total construct thickness of approximately 1.6 mm. The stacked scaffold was maintained under standard culture conditions (37 °C, 5% CO₂) to establish diffusion-limited oxygen and nutrient gradients across the construct. Following incubation, individual paper layers were carefully separated for subsequent RNA extraction and gene expression analysis.

### Doxorubicin treatment of multilayer paper-based 3D tumor cultures

To evaluate the applicability of the optimized RNA extraction workflow for assessing spatially resolved transcriptional responses following drug exposure, multilayer paper-based cultures were treated with doxorubicin (DOX). Following the initial 48 h culture period, the eight-layer constructs (L1–L8) were exposed to complete culture medium containing 5 µM DOX. Untreated multilayer constructs maintained in complete culture medium without DOX served as controls. The constructs were incubated with DOX for 48 h under standard culture conditions (37 °C and 5% CO₂). Following treatment, the multilayer constructs were disassembled, and individual paper layers were collected separately for RNA extraction. RNA extracted from the untreated and DOX-treated layers was subsequently used for q-PCR analysis of *HIF1A* and *VEGFA* expression, with *GAPDH* used as the reference gene.

### RNA extraction from CiGiP using the Qiagen RNeasy kit

MDA-MB-231 cells (1 × 10⁷ cells mL⁻¹) were suspended in Matrigel and seeded onto paper scaffolds to establish cells-in-gel-in-paper (CiGiP) cultures. The scaffolds were placed in 12-well plates and incubated for 48 h before RNA extraction. RNA was extracted using the Qiagen RNeasy Mini Kit (Fig. S1, Supporting Information). RLT lysis buffer was added, followed by homogenization using a Vortex-Genie 2 to facilitate cell disruption and RNA release from the paper–Matrigel matrix. Following centrifugation, the supernatant was transferred to a fresh tube and mixed with 70% ethanol to facilitate RNA binding to the silica membrane. The mixture was transferred to an RNeasy spin column and centrifuged according to the manufacturer’s instructions. The column was washed once with RW1 buffer and twice with RPE buffer before RNA elution with RNase-free water. All centrifugation steps were performed using an Eppendorf 5424R refrigerated microcentrifuge at 8,000–14,000 rpm, as specified in the manufacturer’s protocol.

### RNA extraction from CiGiP using the TRIzol–chloroform method

RNA was extracted from CiGiP scaffolds using a TRIzol–chloroform-based protocol. TRIzol reagent (Thermo Fisher Scientific, Waltham, MA, USA; Cat. No. 15596026) was added directly to the scaffolds, followed by vortex homogenization and incubation at 4 °C. RNA was subsequently isolated through chloroform-mediated phase separation and isopropanol precipitation, followed by ethanol washing and resuspension in RNase-free water.

### Agarose gel for RNA integrity visualization

RNA integrity was assessed using a 1.2% bleach agarose gel. The gel was prepared by dissolving 1.8 g agarose in 150 mL of 1× TBE buffer. The mixture was allowed to stand for 15 min with gentle swirling every 5 min. The suspension was heated until the agarose was completely dissolved and then allowed to cool slightly. SYBR Gold nucleic acid stain was added to the molten agarose according to the manufacturer’s recommended dilution, mixed thoroughly, and the gel was poured into a casting tray and allowed to solidify. RNA samples were mixed with TriTrack RNA loading dye to obtain the appropriate loading concentration and loaded into the gel wells. Electrophoresis was performed in 1× TBE buffer at 100 V for approximately 30 min. RNA bands were visualized using a UV gel documentation system, and RNA integrity was assessed by the presence of distinct 28S and 18S rRNA bands.

### cDNA synthesis

Total RNA was reverse transcribed into complementary DNA (cDNA) using the ProtoScript® II First Strand cDNA Synthesis Kit, New England Biolabs (NEB), Cat. No. E6560S. Prior to reverse transcription, the RNA input was normalized across all samples to 500 ng of total RNA. Each reverse transcription reaction contained 4 µL of 5× LunaScript RT SuperMix and the reaction volume was adjusted to 20 µL with nuclease-free water. Thus, an equivalent amount of total RNA (500 ng) was used for cDNA synthesis from each sample, irrespective of differences in RNA concentration among scaffold layers or experimental conditions. Reverse transcription was performed at 25 °C for 2 min and 55 °C for 10 min, followed by enzyme inactivation at 95 °C for 1 min, according to the manufacturer’s protocol. The synthesized cDNA was stored at −20 °C until further use.

### Quantitative PCR

Quantitative PCR (qPCR) was performed using the Luna® Universal qPCR Master Mix (New England Biolabs, Ipswich, MA, USA) to quantify the expression of *HIF-1A*, *HIF-2A*, *VEGF, GLUT4*, and the housekeeping gene *GAPDH*. The primer sequences used in this study are listed in Table 1. Each qPCR reaction was carried out in a 20 µL reaction volume containing 10 µL of Luna Universal qPCR Master Mix, 0.5 µL each of forward and reverse primers (10 µM), 2 µL of cDNA template, and 7 µL of nuclease-free water. A no-template control (NTC) was included for each primer pair to monitor contamination, and all reactions were performed in triplicate. Thermal cycling was carried out with an initial denaturation at 95°C for 60 s, followed by 40–45 amplification cycles consisting of denaturation at 95°C for 15 s and annealing/extension at 60°C for 30 s, during which fluorescence was acquired at the end of each extension step. Following amplification, a melt curve analysis (60–95°C) was performed to verify the specificity of the amplified products.

**Table 1.**
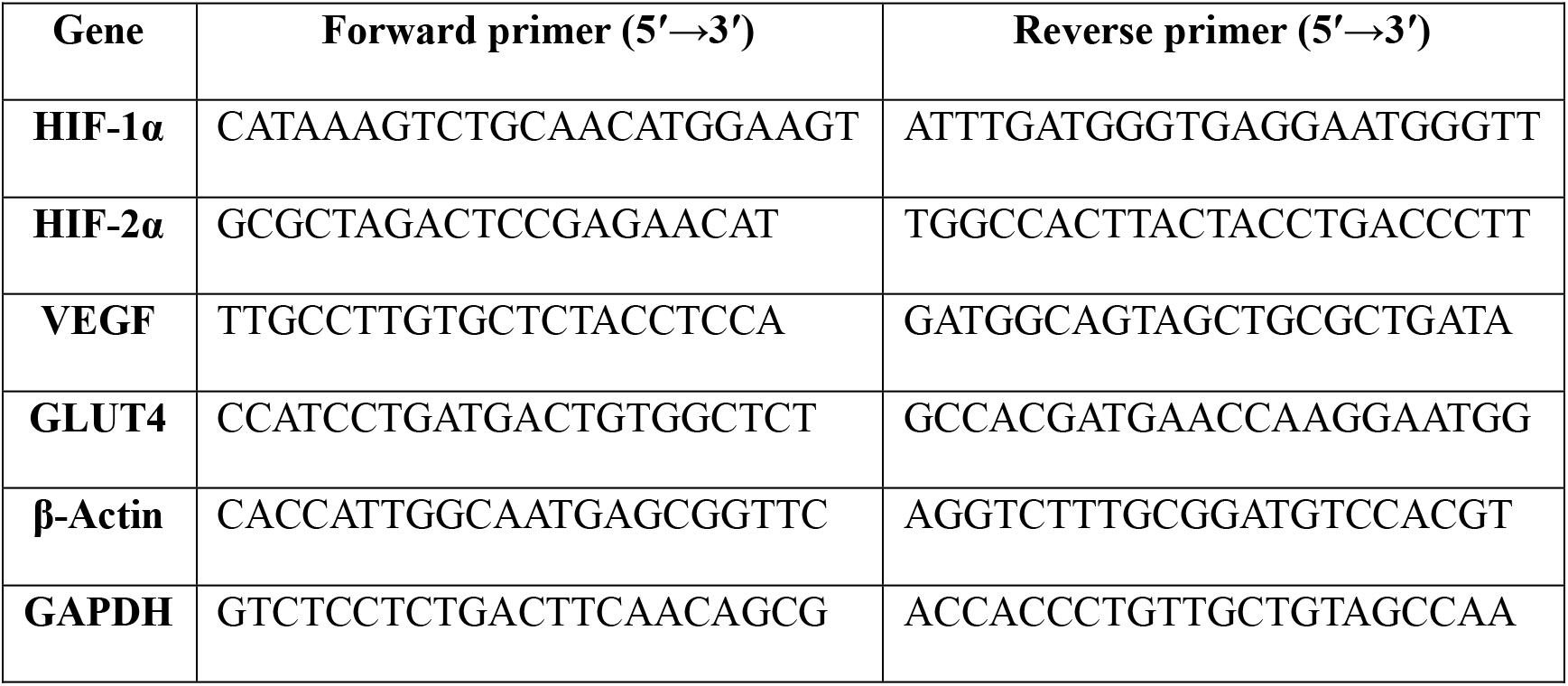
Primer sequences used for quantitative real-time PCR.

### Quantification of layer-resolved relative gene expression

Relative gene expression was quantified using the comparative 2^−ΔΔ*C*^*^t^* method, with *GAPDH* serving as the housekeeping gene for normalization. First, the expression of each target gene was normalized to the corresponding *GAPDH* expression within the same sample to account for variations in RNA input and reverse transcription efficiency. The normalized expression (ΔCt) was calculated using Equation (1)

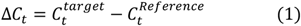

The ΔCt value of each experimental sample was then compared with that of the control sample to determine the ΔΔCt value using Equation (2)

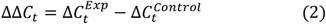

The relative fold change in gene expression was subsequently calculated using Equation (3)

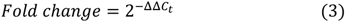

### Statistical Analysis

All experiments were performed using three independent biological replicates (*n* = 3), and data are presented as mean ± standard deviation (SD), unless otherwise stated. Statistical analyses were performed using GraphPad Prism 8.0 (GraphPad Software, San Diego, CA, USA). Comparisons involving multiple groups were performed using one-way analysis of variance (ANOVA) followed by the appropriate multiple-comparisons test, as specified in the corresponding figure legends. For DOX-induced gene-expression analysis, fold-change values for each layer were compared with the theoretical value of 1 using a two-sided one-sample *t*-test. A *p*-value < 0.05 was considered statistically significant.

## Results

### Experimental Framework for Spatially Resolved Gene-Expression Analysis in a Multilayer Paper-Based 3D Tumor Model

To enable spatially resolved molecular characterization of an *in vitro* tumor model, we established an experimental workflow that integrates a multilayer paper-based 3D culture platform with layer-specific RNA extraction and qRT-PCR analysis (Fig. 1). MDA-MB-231 cells embedded in Matrigel were incorporated into individual paper scaffolds, which were assembled into an eight-layer construct (L1–L8) and maintained within a 3D-printed press-fit culture device previously designed by our group^23^. The stacked configuration provides spatially distinct and individually retrievable culture regions while maintaining the cells within a continuous multilayer construct.

**Fig. 1.**
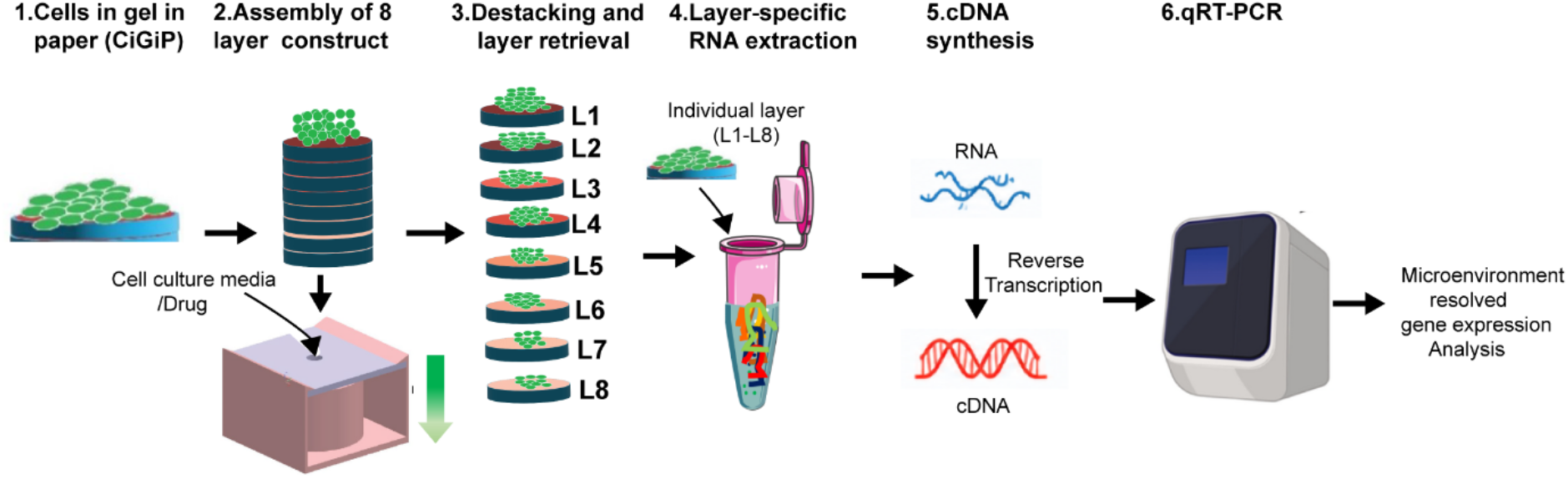
Experimental workflow for spatially resolved gene expression analysis in a multilayer paper-based 3D tumor model. MDA-MB-231 cells embedded in Matrigel were incorporated into individual paper scaffolds and assembled into an eight-layer construct (L1–L8) within a 3D-printed press-fit culture device. Following culture under the specified experimental condition, the construct was disassembled and individual layers were retrieved separately while retaining their spatial identity. RNA was extracted independently from each layer, followed by cDNA synthesis and qPCR analysis, enabling spatially resolved assessment of gene expression across the multilayer construct.

Following culture under the desired experimental condition, including untreated or drug-treated conditions, the construct was disassembled and individual paper layers were retrieved separately while preserving their original spatial positions within the stack. RNA was then extracted independently from each layer, followed by cDNA synthesis and qPCR analysis. This workflow therefore enables gene expression to be interrogated as a function of position within the multilayer construct rather than as a bulk measurement of the entire 3D culture. The subsequent experiments focused on optimizing RNA recovery from the paper–Matrigel matrix and applying the resulting workflow to characterize layer-dependent gene expression and transcriptional responses to doxorubicin treatment.

### Evaluation of silica column-based RNA extraction strategies

Four silica column-based RNA extraction strategies were evaluated for MDA-MB-231 cells cultured under different microenvironmental conditions: conventional Qiagen RNeasy extraction (Fig. 2A), β-mercaptoethanol (βME)-assisted lysis followed by RNeasy purification (Fig. 2B), trypsin pretreatment followed by RNeasy purification (Fig. 2C), and collagenase pretreatment followed by RNeasy purification (Fig. 2D).

**Fig. 2.**
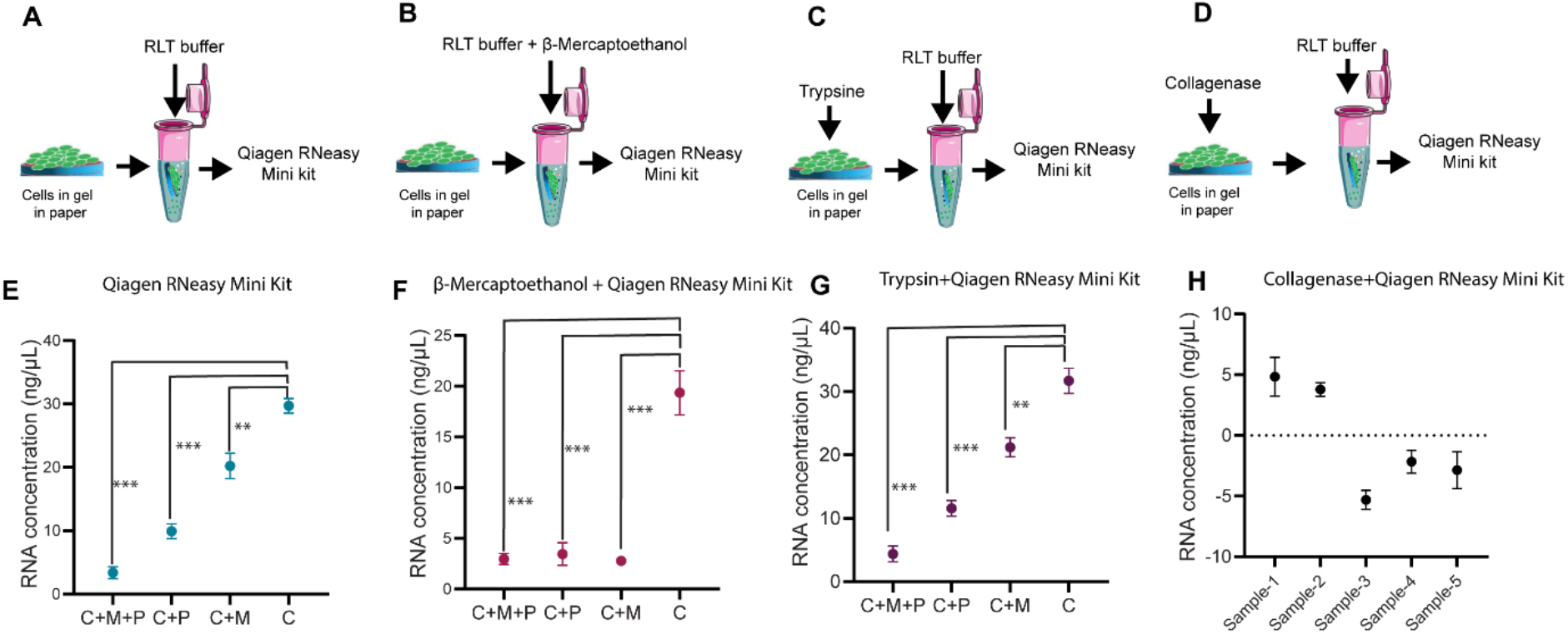
Evaluation of silica column-based RNA extraction strategies across different culture environments. (A, E) Conventional Qiagen RNeasy Mini Kit workflow and the corresponding RNA concentrations obtained from different culture environments. (B, F) β-Mercaptoethanol (βME)-assisted lysis followed by Qiagen RNeasy purification and the corresponding RNA concentrations. (C, G) Trypsin pretreatment followed by Qiagen RNeasy purification and the corresponding RNA concentrations. (D, H) Collagenase pretreatment followed by Qiagen RNeasy purification and the corresponding RNA concentrations. The schematic panels (A–D) illustrate the respective sample-preparation and extraction strategies, whereas panels (E–H) show the resulting RNA recovery across the evaluated culture environments. C, cells cultured on conventional tissue-culture plates; C+M, cells embedded in Matrigel without paper; C+P, cells cultured on paper scaffolds without Matrigel; and C+M+P, cells embedded in Matrigel within paper scaffolds. Data are presented as mean ± SD from three independent biological replicates (n = 3). Statistical significance was assessed using one-way ANOVA followed by Tukey’s multiple-comparisons test. **p < 0.01 and ***p < 0.001.

Prior to evaluating the extraction strategies, the effect of cell density on RNA recovery was examined using MDA-MB-231 cells cultured on conventional tissue culture plates. Increasing the cell density from 10⁶ to 10⁷ cells mL⁻¹ increased the recovered RNA concentration from 14.31 to 154.9 ng µL⁻¹, respectively (Fig. S2 and Table S1, Supporting Information). Based on the substantially higher RNA recovery obtained at the higher cell density, 10⁷ cells mL⁻¹ was selected for subsequent RNA extraction experiments.

Using the conventional Qiagen RNeasy Mini Kit, RNA recovery progressively decreased as the complexity of the cell-culture microenvironment increased (Fig. 2E and Table S2, Supporting Information). Cells cultured directly on conventional tissue culture plates (C) yielded 29.70 ng µL⁻¹ RNA, whereas cells embedded in Matrigel without paper (C+M) yielded 20.21 ng µL⁻¹. Incorporation of cells into paper scaffolds without Matrigel (C+P) further reduced the recovered RNA concentration to 9.91 ng µL⁻¹. The lowest RNA recovery, 3.39 ng µL⁻¹, was obtained when cells were embedded in Matrigel within the paper scaffold (C+M+P). Thus, the simultaneous presence of the cellulose scaffold and Matrigel represented the most challenging condition for RNA recovery using the conventional silica column-based method.

To determine whether enhanced lysis could improve RNA recovery from the scaffold, βME was incorporated into the extraction procedure (Fig. 2B). However, βME-assisted lysis produced a similar dependence of RNA recovery on culture configuration (Fig. 2F and Table S3, Supporting Information). RNA concentrations of 31.70, 21.21, 11.58, and 4.39 ng µL⁻¹ were obtained from C, C+M, C+P, and C+M+P samples, respectively. Although βME-assisted lysis resulted in a small increase in RNA recovery from the C+M+P condition compared with the conventional Qiagen procedure, the recovered RNA concentration remained substantially lower than that obtained from cells cultured without the combined paper–Matrigel matrix.

Enzymatic pretreatment was subsequently investigated to determine whether disruption of the cellular and matrix environment before silica column purification could improve RNA recovery. Trypsin pretreatment (Fig. 2C) did not improve RNA recovery from the paper–Matrigel construct (Fig. 2G and Table S4, Supporting Information). RNA concentrations of 19.36, 2.78, 3.45, and 2.97 ng µL⁻¹ were obtained from C, C+M, C+P, and C+M+P samples, respectively. In particular, the C+M+P condition yielded less than 3 ng µL⁻¹ RNA despite trypsin pretreatment, indicating that enzymatic cell detachment prior to column purification was insufficient to overcome the limitations associated with RNA recovery from the composite scaffold. Collagenase pretreatment was also evaluated as an alternative strategy for disrupting the matrix prior to RNA isolation (Fig. 2D); however, this approach provided very poor recovery even for cells grown on plates (Fig. 2H; Table S5, Supporting Information), so more complex microenvironments involving paper and matrix were not tried with this method.

For the Qiagen, βME, and trypsin-based extraction strategies, one-way ANOVA demonstrated significant differences in RNA concentration among the investigated culture conditions (p < 0.0001; Fig. 2E–G). Across these extraction strategies, RNA recovery was consistently highest from cells cultured on conventional tissue culture plates and markedly reduced when cells were incorporated into scaffold-based culture environments. Most importantly, the combined C+M+P condition consistently produced the lowest or among the lowest RNA concentrations, demonstrating that conventional silica column-based extraction, even when supplemented with chemical or enzymatic pretreatment, was insufficient for efficient RNA recovery from the paper–Matrigel 3D culture system. These findings motivated the subsequent evaluation of alternative extraction approaches specifically aimed at improving RNA recovery from the composite scaffold.

### Optimization and selection of TRIzol–chloroform extraction for the CiGiP

Since the silica column-based extraction strategies resulted in limited RNA recovery from cells embedded within the paper–Matrigel scaffold, a TRIzol–chloroform extraction procedure was subsequently evaluated. The optimized workflow is illustrated in Fig. 3A. Briefly, the CiGiP scaffold was incubated directly with TRIzol reagent, followed by vortexing and centrifugation to separate the paper scaffold and insoluble debris from the lysate. The recovered TRIzol lysate was subjected to chloroform-mediated phase separation, and the RNA-containing aqueous phase was collected. RNA was subsequently precipitated using isopropanol, washed with 70% ethanol, air-dried, resuspended in RNase-free water, and quantified spectrophotometrically.

**Fig. 3.**
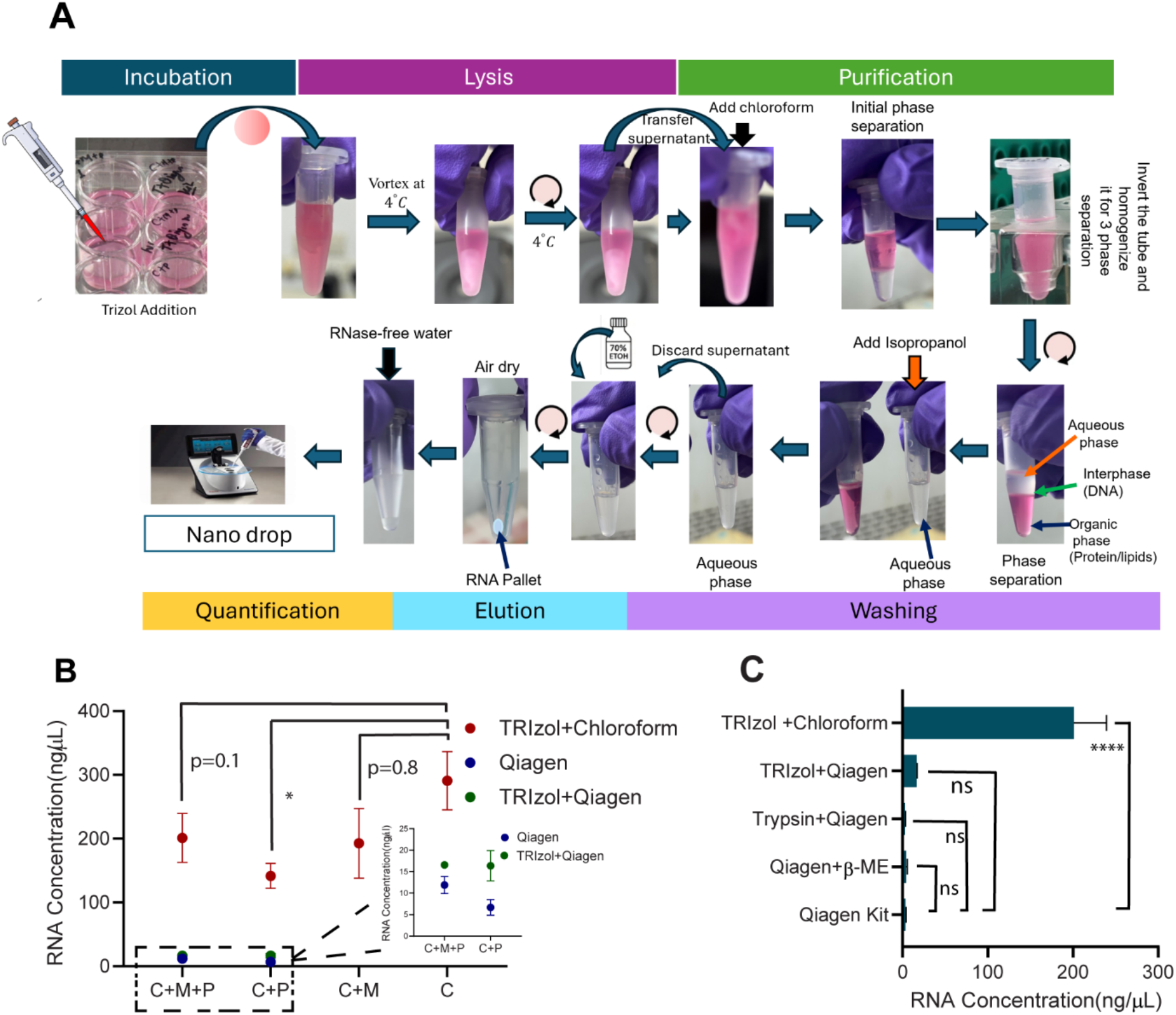
Optimization and selection of the TRIzol–chloroform method for RNA extraction from paper-based 3D cultures. (A) Experimental workflow for TRIzol–chloroform extraction from CiGiP, comprising TRIzol-mediated lysis, removal of the paper scaffold and insoluble debris, chloroform-mediated phase separation, recovery of the aqueous phase, isopropanol precipitation, ethanol washing, RNA resuspension, and spectrophotometric quantification. (B) RNA concentrations obtained using the TRIzol–chloroform method across different culture microenvironments. The inset compares RNA recovery using the Qiagen RNeasy Mini Kit and the combined TRIzol–Qiagen workflow for scaffold-containing conditions. (C) Direct comparison of RNA recovery from cells embedded in the paper–Matrigel scaffold (C+M+P) using the Qiagen RNeasy Mini Kit, β-Mercaptoethanol-assisted Qiagen extraction, trypsin pretreatment followed by Qiagen extraction, the combined TRIzol–Qiagen workflow, and conventional TRIzol–chloroform extraction. C, cells cultured on conventional tissue culture plates; C+M, cells embedded in Matrigel; C+P, cells cultured on paper scaffolds; and C+M+P, cells embedded in Matrigel within paper scaffolds. Data are presented as mean ± SD (n = 3 independent biological replicates). Statistical significance was determined using one-way ANOVA followed by Tukey’s multiple-comparisons test for panel B and one-way ANOVA followed by Dunnett’s multiple-comparisons test for panel C, using the Qiagen RNeasy Mini Kit as the reference group. ns, not significant; *p < 0.05; ****p < 0.0001.

The TRIzol–chloroform procedure substantially improved RNA recovery across the investigated culture microenvironments (Fig. 3B). Mean RNA concentrations of approximately 201, 141, 192, and 290 ng µL⁻¹ were obtained from cells cultured in paper–Matrigel (C+M+P), paper alone (C+P), Matrigel alone (C+M), and conventional tissue culture plates (C), respectively. Unlike the silica column-based approaches, the TRIzol–chloroform procedure therefore enabled recovery of high RNA concentrations even from the combined paper–Matrigel scaffold. Although the overall RNA concentration differed among the culture configurations, RNA recovery from the C+M+P condition was not significantly different from that obtained from C+M (p = 0.8) or conventional tissue culture conditions (p = 0.1), whereas a significant difference was observed between C+M+P and C+P (Fig. 3B).

To determine whether the improved recovery was attributable primarily to TRIzol-mediated lysis or chloroform extraction, a combined TRIzol–Qiagen workflow was also evaluated. In this approach, TRIzol was used for initial lysis, followed by silica column-based purification. Incorporation of TRIzol prior to Qiagen purification increased RNA recovery compared with the conventional Qiagen procedure for scaffold-containing samples (inset, Fig. 3B). However, RNA concentrations obtained using the combined TRIzol–Qiagen workflow remained markedly lower than those obtained using the complete TRIzol–chloroform extraction procedure. This observation indicates that improved lysis alone did not account for the high RNA recovery and that the subsequent phase-separation and precipitation steps contributed substantially to efficient RNA isolation from the composite scaffold.

A direct comparison of the extraction strategies was subsequently performed using the C+M+P condition, which represented the most challenging configuration for RNA recovery (Fig. 3C). The conventional Qiagen RNeasy procedure yielded 3.39 ng µL⁻¹ RNA. Neither βME-assisted Qiagen extraction (4.39 ng µL⁻¹) nor trypsin pretreatment followed by Qiagen purification (2.97 ng µL⁻¹) significantly improved RNA recovery relative to the conventional Qiagen procedure. The combined TRIzol–Qiagen workflow increased the RNA concentration to 16.58 ng µL⁻¹; however, this increase was not statistically significant relative to the Qiagen reference. The TRIzol–chloroform procedure yielded 201.13 ng µL⁻¹ RNA and was significantly higher than the conventional Qiagen extraction (p < 0.0001). This corresponded to an approximately 59-fold increase in recovered RNA concentration relative to the conventional Qiagen procedure.

Representative agarose gel electrophoresis images were used to further assess the recovered RNA and are provided in Fig. S3 (Supporting Information). RNA obtained using the conventional Qiagen-based procedure produced comparatively weak rRNA bands, particularly for scaffold-containing samples, whereas TRIzol–chloroform extraction produced substantially stronger and readily detectable 28S and 18S rRNA bands. The combined TRIzol–Qiagen workflow produced detectable RNA but remained inferior to the complete TRIzol–chloroform procedure, consistent with the quantitative RNA concentration measurements. Collectively, these results identified the TRIzol–chloroform procedure as the most effective extraction strategy investigated for recovering RNA from cells embedded within the paper–Matrigel scaffold. The TRIzol–chloroform method was therefore selected for subsequent layer-specific RNA extraction and gene-expression analysis.

### RNA extraction from multilayer CiGiP culture

Following optimization of the RNA extraction procedure, the ability of the optimized TRIzol–chloroform method to recover RNA from spatially distinct regions of the multilayer paper-based 3D culture system was evaluated. The 3D-printed press-fit culture device accommodated eight stacked paper layers (L1–L8), which could be individually retrieved following culture and processed independently for RNA extraction (Fig. 4A). This configuration enabled preservation of the spatial identity of each layer throughout RNA isolation and subsequent molecular analysis. Quantifiable RNA was successfully recovered from all eight layers of the multilayer construct (Fig. 4A and Table S6, Supporting Information). A single-layer (SL) culture (not stacked), used as a control, yielded an RNA concentration of 288.53 ng µL⁻¹. Within the multilayer construct, the highest RNA concentration was observed in L1 (245.26 ng µL⁻¹), followed by a general trend of decrease with increasing scaffold depth. RNA concentrations across L1–L8 ranged from 113.06 to 245.26 ng µL⁻¹, demonstrating measurable RNA recovery throughout the entire eight-layer construct (Fig. 4A). Although some layer-to-layer variation was observed, including a modest increase in L7 relative to L6, substantial quantities of RNA were recovered even from the deepest layers of the construct.

**Fig. 4.**
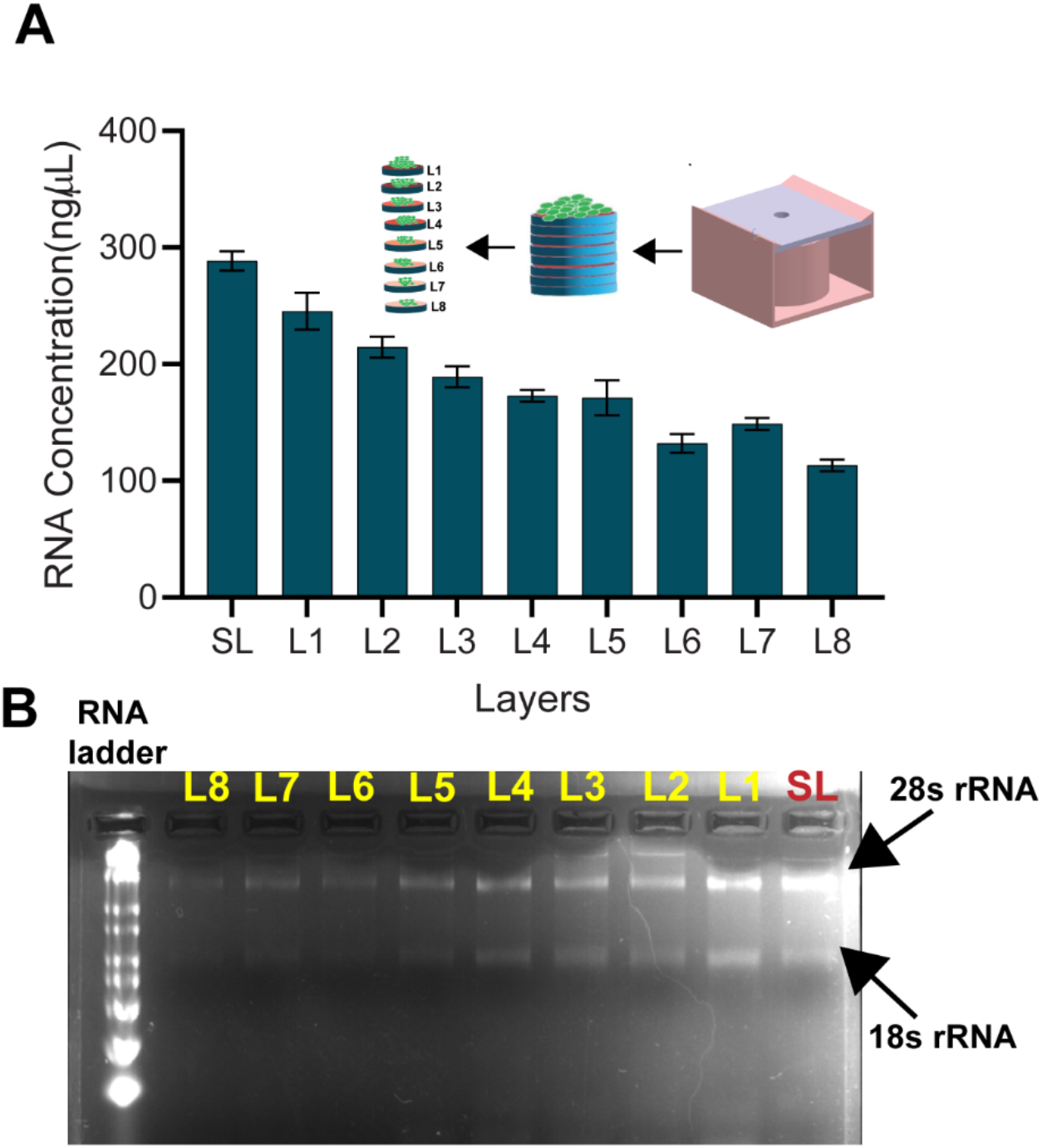
Layer-specific RNA recovery and integrity assessment from the multilayer paper-based 3D culture system. (A) RNA concentration recovered from individual scaffold layers using the optimized TRIzol–chloroform extraction protocol. Data are presented as mean ± SD (n = 3 independent biological replicates). (B) Representative agarose gel electrophoresis image of RNA isolated from individual scaffold layers, showing the RNA ladder and the 28S and 18S rRNA regions used for assessment of RNA integrity. SL represents cells grown on single layer

In addition to RNA concentration, purity was assessed using the A260/A280 and A260/A230 absorbance ratios (Table S6, Supporting Information). The recovered RNA across all layers exhibited A260/A280 ratios ranging from 1.78 to 1.96 and A260/A230 ratios ranging from 1.93 to 2.14. Ideal reference ranges for these ratios for high purity RNA are < 2 and < 2.2, respectively, supporting the suitability of the extracted RNA for subsequent molecular analysis.

RNA integrity following layer-specific extraction was further assessed by agarose gel electrophoresis (Fig. 4B). Ribosomal RNA bands corresponding to the 28S and 18S rRNA regions were detectable in RNA recovered from the individual scaffold layers, although band intensity varied across the stack. Together, the RNA concentration, purity, and electrophoretic assessment demonstrate that the optimized TRIzol–chloroform procedure enabled recovery of measurable and sufficiently intact RNA from individual spatial locations throughout the multilayer construct, thereby permitting subsequent layer-specific gene expression analysis.

To further evaluate the applicability of the extraction workflow under a pharmacological perturbation, RNA was also recovered from individual layers following doxorubicin treatment. Quantifiable RNA was obtained from all L1–L8 layers under both untreated and DOX-treated conditions (Fig. S4A and Table S7 Supporting Information). Although DOX exposure altered the amount of RNA recovered from several layers, sufficient RNA was obtained across the construct for subsequent cDNA synthesis and qRT-PCR analysis. Representative agarose gel electrophoresis further confirmed the recovery of RNA from both untreated and DOX-treated layers (Fig. S4B-C, Supporting Information). These results demonstrate that the optimized workflow applies not only to untreated but also to drug-exposed multilayer cultures.

### Spatially resolved gene expression in untreated multilayer tumor cultures

Following validation of layer-specific RNA recovery, the optimized workflow was applied to investigate spatial variations in gene expression across the multilayer paper-based 3D tumor model. Representative Calcein-AM fluorescence images showed a progressive reduction in viable-cell fluorescence from the upper toward the deeper scaffold layers (Fig. 5A). Quantitative analysis similarly demonstrated a depth-dependent decrease in cell viability relative to the single layer control (SL; Fig. 5B). Cell viability decreased from approximately 81% of SL in L1 to 22–26% of SL in the deeper layers (L5–L8), where the response approached a plateau. One-way ANOVA demonstrated a significant overall effect of scaffold layer on cell viability (p < 0.0001). Dunnett’s multiple comparisons test further showed significantly reduced viability in the scaffold layers L4-L8 relative to SL control, with the magnitude of reduction generally increasing with scaffold depth (Table S9, Supporting Information). These observations are consistent with the development of increasingly restrictive microenvironmental conditions toward the interior of the multilayer construct.

**Fig. 5.**
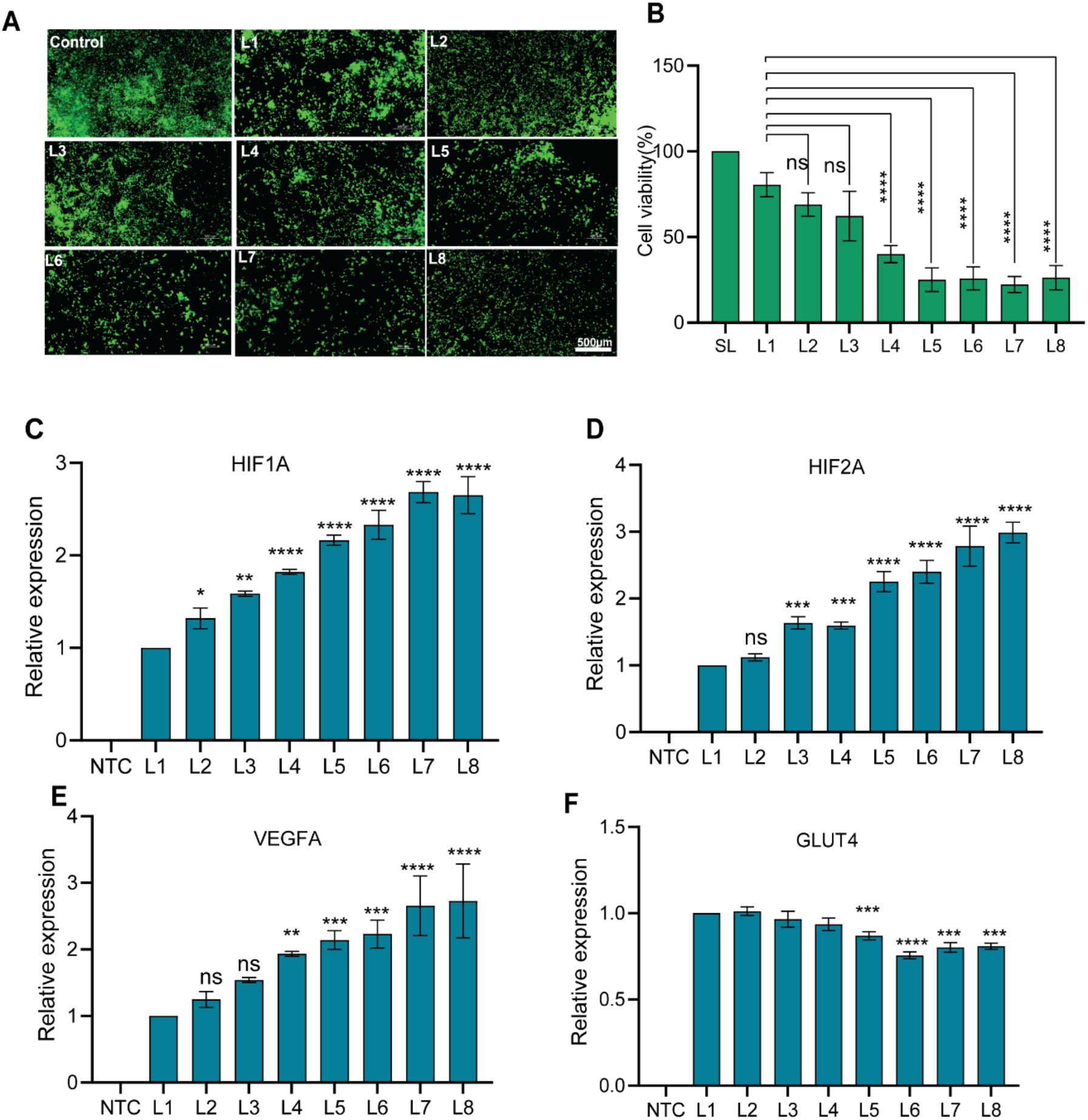
Spatially resolved cellular viability and gene expression across the multilayer paper-based 3D tumor model. (A) Representative Calcein-AM fluorescence images showing viable-cell distribution across scaffold layers L1–L8 compared with L1. (B) Quantification of cell viability across scaffold layers, expressed relative to the 2D control. (C–F) Relative expression of (C) HIF-1A, (D) HIF-2A, (E) VEGFA, and (F) GLUT4 across scaffold layers determined by qRT-PCR. Target-gene expression was normalized to GAPDH and calculated using the 2^−ΔΔCt^ method with L1 as the calibrator. Data are presented as mean ± SD (n = 3 independent biological replicates). Statistical significance was evaluated using the corresponding ANOVA followed by Dunnett’s multiple-comparisons test. *p < 0.05, **p < 0.01, ***p < 0.001, ****p < 0.0001. Scale bar = 500 µm.

Layer-specific transcriptional responses were subsequently evaluated by qPCR *for HIF-1A, HIF-2A, VEGFA*, and *GLUT4* (Fig. 5C–F), using GAPDH as a housekeeping gene. Prior to performing transcriptional studies, primer amplification efficiencies were validated using standard curves generated from serial cDNA dilutions (Fig. S5, Supplementary Information). The calculated amplification efficiencies are summarized in Table S7 (Supplementary Information). The relative gene expression across the multilayer construct was calculated using the 2^−ΔΔ*C*^*^t^* method with L1 as the calibrator. *HIF-1A* expression exhibited an overall increase toward the deeper scaffold layers, with the strongest expression observed in the lower regions of the construct (Fig. 5C and Table S10, Supporting Information). *HIF-2A* displayed a similar depth-dependent response, with increased expression in the deeper layers relative to L1 (Fig. 5D and Table S11, Supporting Information). These changes indicate that the spatially heterogeneous microenvironment established within the multilayer construct was accompanied by differential regulation of hypoxia-associated genes. *VEGF* expression also increased with scaffold depth (Fig. 5E and Table S12, Supporting Information), consistent with the observed changes in *HIF*-associated transcriptional responses. In contrast, *GLUT4* exhibited an overall decrease toward the deeper scaffold layers (Fig. 5F and Table S13, Supporting Information), indicating that different molecular pathways responded distinctly to the spatial microenvironment within the construct. Statistical analysis further supported layer-dependent differences in gene expression. One-way repeated-measures analysis demonstrated significant variation in expression across the scaffold layers, followed by Dunnett’s multiple-comparisons test using L1 as the reference layer. The corresponding adjusted p-values for *HIF-1A, HIF-2A, VEGFA*, and *GLUT4* are provided in Tables S10–S13 (Supporting Information).

These molecular findings complement our previous observations of depth-dependent cellular responses within the same multilayer scaffold platform. Importantly, the present analysis extends those observations by demonstrating that individual scaffold layers can be independently recovered and interrogated at the transcriptional level. Collectively, the viability and qRT-PCR results demonstrate the utility of the optimized RNA extraction workflow for spatially resolved molecular characterization within the multilayer paper-based *in vitro* tumor model.

### Spatially resolved transcriptional response to doxorubicin treatment

To evaluate spatially dependent responses to pharmacological perturbation, multilayer paper-based tumor cultures were exposed to 5 µM doxorubicin (DOX) for 48 h. Cell viability was first evaluated independently at each scaffold position and expressed relative to the corresponding layer of an untreated multilayer culture, thereby accounting for the pre-existing layer-dependent differences in cell viability within the construct. Thus, viability in DOX-treated L1 was normalized to untreated L1, DOX-treated L2 to untreated L2, and similarly through L8. DOX treatment produced a pronounced but non-monotonic spatial variation in relative cell viability (Fig. 6A). Viability was lowest in the upper layers, with approximately 25–32% of the corresponding untreated control in L1–L3, increased to approximately 51% in L4, and reached a maximum of approximately 67% in L5. Relative viability subsequently decreased to approximately 53% in L6, 34% in L7, and 26% in L8. These results demonstrate that the phenotypic response to DOX was strongly dependent on spatial position within the multilayer construct, and point to the development of a drug-resistant region around the intermediate layers L4-L6.

**Fig. 6.**
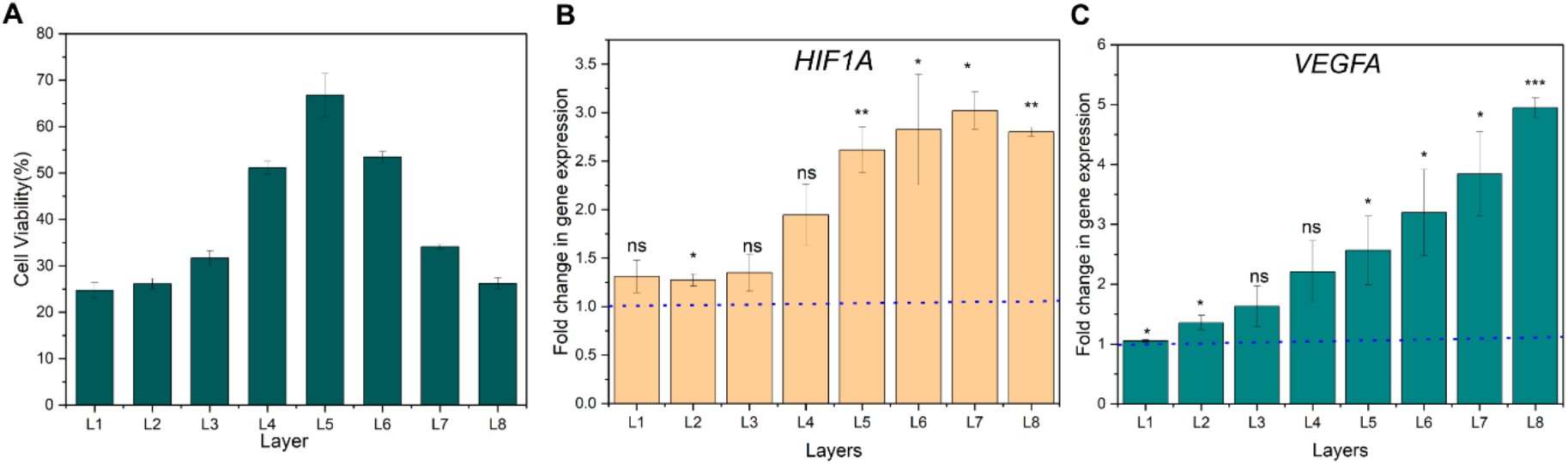
Layer-resolved transcriptional response to doxorubicin treatment in the multilayer paper-based 3D tumor model. (A) Cell viability across scaffold layers L1–L8 following DOX treatment, expressed relative to the corresponding untreated layer. Relative expression of (B) HIF-1A and (C) VEGFA across scaffold layers L1–L8 following DOX treatment. For each scaffold position, expression in the DOX-treated sample was normalized to the corresponding untreated control using the 2^−ΔΔCt^ method after normalization to GAPDH. The horizontal dashed line at a fold change of 1 represents no change relative to the corresponding untreated control. Data are presented as mean ± SD from three independent biological replicates (n = 3). Statistical significance at each layer was determined using a two-sided one-sample t-test comparing the fold-change values with the hypothetical value of 1. *p < 0.05, **p < 0.01, ***p < 0.001; ns, not significant.

To distinguish DOX-induced transcriptional changes from the pre-existing depth-dependent differences observed in untreated cultures, gene expression in each DOX-treated layer was compared with its corresponding untreated layer. Consequently, a fold change of 1 represents no change in gene expression following DOX treatment at the corresponding scaffold position. DOX treatment produced a spatially heterogeneous *HIF-1A* transcriptional response across the multilayer construct (Fig. 6B). The mean HIF-1A fold change increased from 1.42-fold in L1 to approximately 2.7–2.9-fold in L5–L8. Compared with the no-change reference value of 1, *HIF-1A* expression was significantly increased in L2 (p = 0.038), L5 (p = 0.0049), L6 (p = 0.0295), L7 (p = 0.0369), and L8 (p = 0.0052), whereas the changes observed in L1, L3, and L4 were not statistically significant (Table S14, Supporting Information). The greater fold changes observed in the deeper regions of the construct indicate that the transcriptional response to DOX varied according to spatial position within the multilayer tumor model. *VEGFA* exhibited a more pronounced depth-associated response to DOX (Fig. 6C). Mean *VEGFA* expression increased from approximately 1.05-fold in L1 to 4.95-fold in L8. Significant increases relative to the no-change reference were observed in L1 (p = 0.0356), L2 (p = 0.0367), L5 (p = 0.0419), L6 (p = 0.0339), L7 (p = 0.0200), and L8 (p = 0.00057), whereas changes in L3 and L4 were not statistically significant (Table S14, Supporting Information). Notably, the magnitude of VEGFA induction increased substantially toward the deeper scaffold layers, with the strongest response observed in L8. Note that melt-curve analysis was performed to assess qRT-PCR product specificity for *GAPDH*, *HIF-1A*, and *VEGFA* in untreated and DOX-treated samples (Figs. S6 and S7, Supporting Information).

Together, these results demonstrate that the developed workflow can resolve not only baseline spatial differences in gene expression within the multilayer construct but also position-dependent transcriptional responses following drug exposure. This extends the analytical utility of the platform from characterization of intrinsic microenvironmental heterogeneity to spatial assessment of drug-induced molecular responses.

## Discussion

Efficient RNA recovery from the composite paper–Matrigel scaffold is essential for molecular analysis of CiGiP cultures. Direct exposure of the scaffold to TRIzol, combined with vortex homogenization, facilitated physical disruption of the paper–Matrigel construct and release of the embedded cellular contents. The improved RNA recovery may be attributed to phenol–guanidinium-mediated cell lysis and RNase inactivation, followed by chloroform-induced phase separation, which facilitates RNA isolation from DNA, proteins, and other cellular components^24^. Importantly, the recovered RNA was suitable for qRT-PCR analysis of individually retrieved scaffold layers, overcoming a key sample-preparation challenge in spatially resolved molecular analysis.

The layer-dependent expression of *HIF1A*, *HIF2A*, and *VEGFA* may reflect heterogeneous cellular responses to the local microenvironment. HIF signalling regulates responses to reduced oxygen availability and contributes to processes such as metabolic regulation and angiogenesis, including *VEGFA* expression. Following DOX treatment, the generally greater induction of *HIF1A* and *VEGFA* in deeper layers contrasted with the non-monotonic viability response, suggesting that drug-induced transcriptional changes cannot be inferred from viability measurements alone. To the best of our knowledge, the present study provides an initial demonstration of layer-resolved DOX-associated *HIF1A* and *VEGFA* induction in a multilayer CiGiP tumor model.

The decreasing *GLUT4* expression toward deeper layers contrasted with the HIF-associated genes, potentially reflecting transporter-specific regulation, as GLUT1 and GLUT3 are more closely associated with HIF signaling^25^. Previous studies have reported distinct GLUT transporter expression patterns in breast cancer and demonstrated the role of GLUT4 in glucose uptake and metabolism in MDA-MB-231 cells^26^. However, the mechanism underlying the observed depth-dependent decrease in *GLUT4* remains unclear.

The ability to recover RNA from physically addressable layers could provide drug developers with molecular information that is lost during bulk analysis of 3D cultures. By combining layer-specific viability measurements with transcriptional profiling, the platform may help identify regions exhibiting distinct responses to therapeutic exposure, including molecular changes that do not directly parallel cell survival. Such information could support mechanistic studies of spatially heterogeneous drug responses and guide the selection of additional molecular markers or treatment conditions for investigation.

Several limitations should be considered. Spatial resolution is restricted to individual scaffold layers, and the targeted qRT-PCR panel captures only a limited range of transcriptional responses. Oxygen and nutrient concentrations, metabolic activity, and local DOX exposure were not directly measured. Future integration of microenvironmental measurements, drug-distribution analysis, and broader molecular profiling would help establish the mechanisms underlying the observed spatial responses.

## Conclusion

This study establishes an effective TRIzol–chloroform-based workflow for RNA recovery from multilayer paper-based 3D tumor cultures. The approach enables layer-specific gene-expression analysis while preserving spatial information. Its application revealed heterogeneous expression of *HIF1A*, *HIF2A*, *VEGFA*, and *GLUT4* across the scaffold layers. Following DOX treatment, transcriptional responses varied with scaffold depth and did not consistently parallel cell viability. These findings highlight the importance of integrating molecular and phenotypic measurements to characterize spatially heterogeneous drug responses. The developed platform offers an accessible approach for investigating tumor-associated transcriptional heterogeneity. Its ability to identify localized molecular responses may also support the evaluation of anticancer therapeutics in physiologically relevant 3D culture models.

## Author Contribution

A.K: methodology, investigation, formal analysis, writing – original draft; B.J.T: conceptualization, methodology, writing – review & editing, supervision, funding acquisition.

## Acknowledgements

This work was supported primarily by a CSR grant from Syngene International Ltd. Additional support was provided by an extramural research grant (BT/PR50679/MED/32/998/2023) from the Department of Biotechnology, India; a core research grant (CRG/2021/003397) from the Science and Engineering Research Board, India; and a research grant by the Isaac Center for Public Health (ICPH) at the Indian Institute of Science.

## Data availability

The data supporting this article are included in the main manuscript and Supporting Information (SI). The Supporting Information contains additional details on RNA extraction and optimization, RNA concentration and purity measurements, primer efficiency analysis, standard curves, and supporting qRT-PCR characterization of the multilayer paper-based 3D tumor model.

## Conflict of Interest Statement

The authors declare no competing financial interests.

## Notes

### Competing Interest Statement

The authors have declared no competing interest.

