## Supplementary Information for "Enabling Spatially Resolved Gene Expression Analysis in a Multilayer Paper-Based *In Vitro* Tumor Model"

[2]Department of Bioengineering
Indian Institute of Science
Bengaluru, Karnataka 560012
India

Keywords: *paper-based 3D culture; tumor microenvironment; spatially resolved gene expression; RNA extraction; qRT-PCR; doxorubicin; MDA-MB-231;CiGiP*

*Correspondence to:
Bhushan J. Toley
Department of Chemical Engineering
Indian Institute of Science, Bangalore
Malleswaram
Bangalore 560012


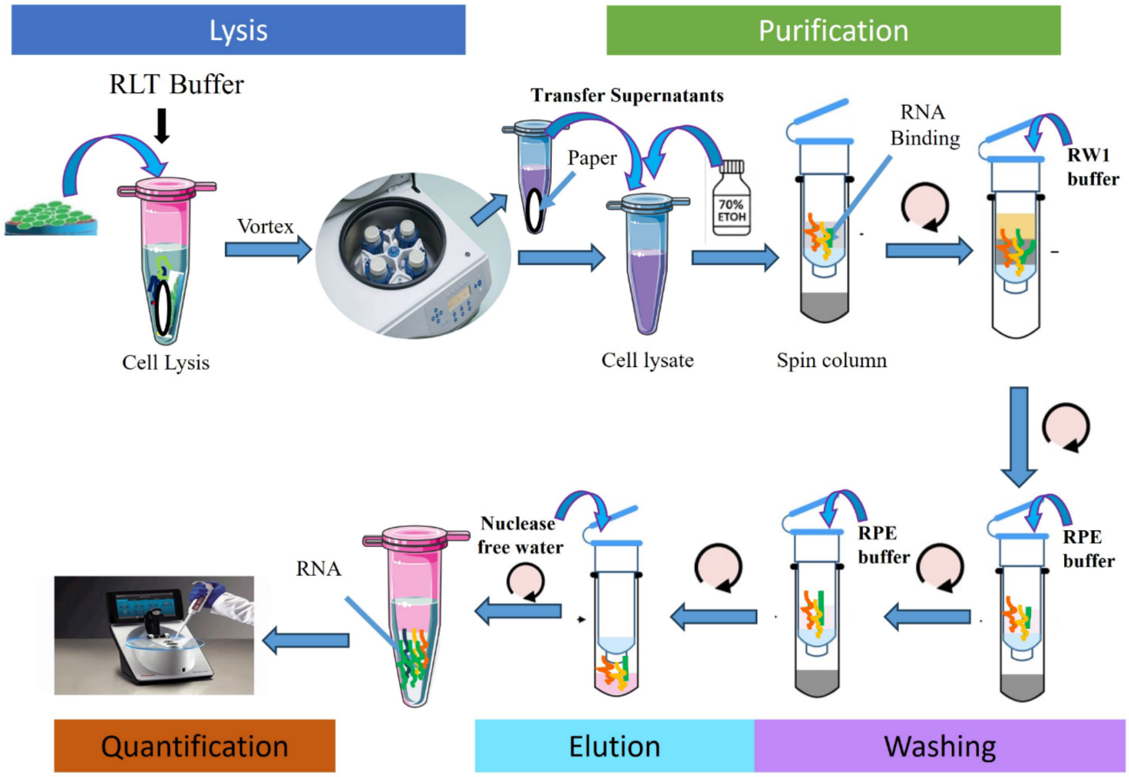


**Fig.S1 .** *Schematic representation of the RNA extraction workflow using the Qiagen RNeasy Mini Kit*

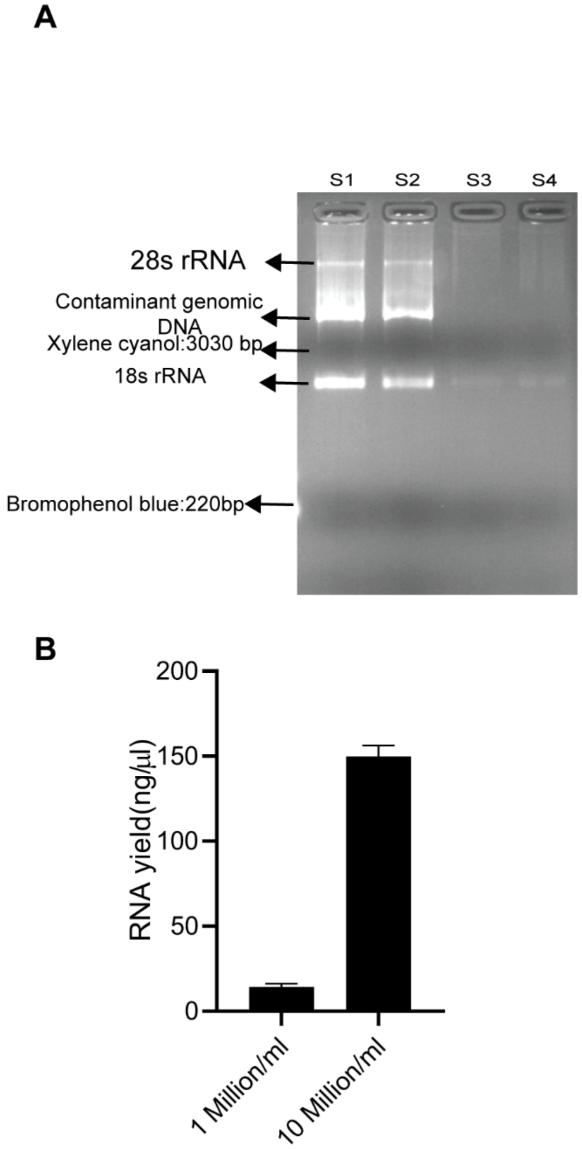


***Fig. S2. Optimization of cell density for RNA extraction.***
*(A) Representative agarose gel electrophoresis image of total RNA extracted from MDA-MB-231 cells cultured on standard 12-well plates at densities of $1 \times 10^6$ cells/mL (S3, S4) and $1 \times 10^7$ cells/mL (S1, S2). (B) Comparison of RNA yield obtained from the two cell densities, demonstrating increased RNA recovery at higher cell density.*

**Table S1.**RNA concentration and purity of total RNA extracted from MDA-MB-231 cells cultured on standard tissue culture plates at two different cell densities.

| Sample ID | Cell Density | RNA concentration (ng/µL) | A260/A280 | A260/A230 |
|---|---|---|---|---|
| Sample 1 | 1 million | 14.31±1.14 | 2.03 | 1.95 |
| Sample 2 | 10 million | 154.9±3.36 | 2.08 | 2.08 |

**Table S2.** RNA concentration and purity (A260/A280 and A260/A230 ratios) of samples extracted using the Qiagen RNeasy Mini Kit.

| **Samples** | **Mean RNA concentration (ng/µl)** | **A260/A280** | **A260/A230** |
|---|---|---|---|
| Cells without paper | 29.7±1.14 | 2.19 | 0.23 |
| Cells in Matrigel without paper | 20.21±1.99 | 2.10 | 0.44 |
| Cells in paper | 9.91±1.17 | 2.39 | 0.06 |
| Cells in Matrigel in paper | 3.39±0.93 | 2.24 | 0.11 |

**Table S3.**RNA concentration and purity (A260/A280 and A260/A230 ratios) of samples extracted following β-mercaptoethanol-assisted lysis and Qiagen RNeasy Mini Kit extraction.

| **Samples** | **Mean RNA concentration (ng/µl)** | **A260/A280** | **A260/A230** |
|---|---|---|---|
| Cells without paper | 31.7±1.97 | 2.19 | 0.43 |
| Cells in Matrigel without paper | 21.213±1.41 | 2.10 | 1.51 |
| Cells in paper | 11.583±1.23 | 2.39 | 1.12 |
| Cells in Matrigel in paper | 4.39±1.25 | 2.24 | 0.04 |

**Table S4.**RNA concentration and purity (A260/A280 and A260/A230 ratios) of samples extracted following trypsin pre-treatment and Qiagen RNeasy Mini Kit extraction.

| **Samples** | **Mean RNA concentration (ng/µl)** | **A260/A280** | **A260/A230** |
|---|---|---|---|
| Cells without paper | 19.36±2.17 | 2.06 | 1.63 |
| Cells in Matrigel without paper | 2.78±0.36 | 2.13 | 1.24 |
| Cells in paper | 3.45±1.11 | 2.32 | 0.06 |
| Cells in Matrigel in paper | 2.97±0.54 | 2.15 | 1.66 |

**Table S5.**RNA concentration and purity (A260/A280 and A260/A230 ratios) of samples extracted following collagenase pre-treatment and Qiagen RNeasy Mini Kit extraction.

| Samples | Mean RNA concentrat (ng/µl) | A260/A280 | A260/A230 |
|---|---|---|---|
| S1 | 4.82±1.60 | 2.39 | 0.13 |
| S2 | 3.785±0.55 | 2.20 | 1.28 |
| S3 | ND | ND | ND |
| S4 | ND | ND | ND |
| S5 | ND | ND | ND |

**ND:** RNA concentration below the detection limit of the NanoDrop spectrophotometer; the instrument returned a negative value after blank correction.

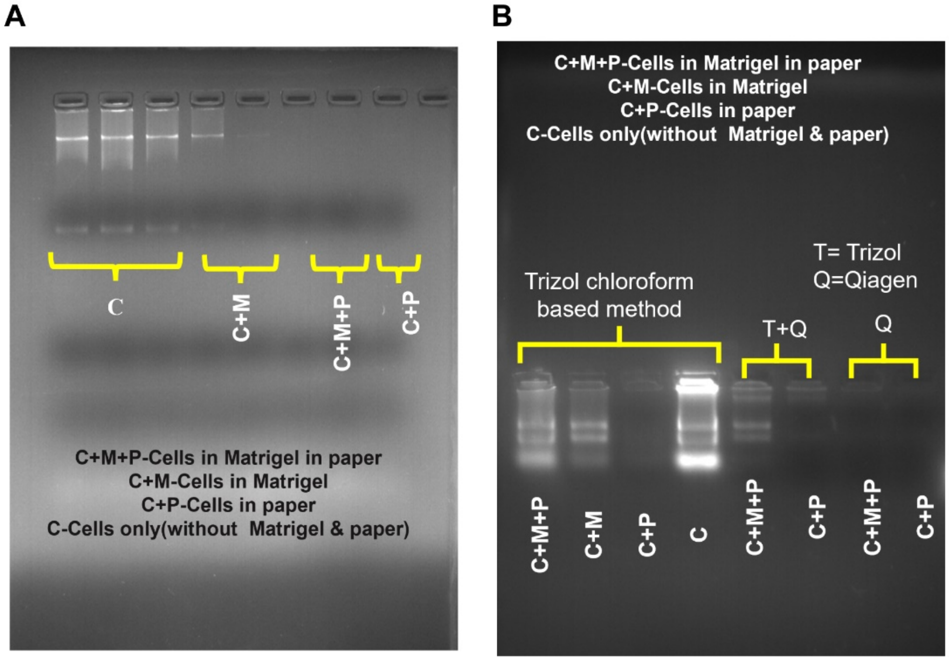


***Fig.S3*** *(A) Agarose gel electrophoresis of RNA extracted from cells cultured under different microenvironmental conditions using the Qiagen-based protocol. (B) Comparative agarose gel electrophoresis of RNA obtained using TRIzol–chloroform extraction, TRIzol followed by Qiagen purification, and the Qiagen kit. C, cells cultured on tissue-culture plastic; C+M, cells embedded in Matrigel; C+P, cells cultured in paper; C+M+P, cells embedded in Matrigel within paper.*

**Table S6.** RNA concentration and purity (A260/A280 and A260/A230 ratios) of RNA extracted from individual layers of the multilayer paper scaffold using the Trizol–chloroform method.

| S.N | Samples | Mean RNA Concentration (ng/µl) | A260/A280 | A260/A230 |
|---|---|---|---|---|
| 1 | SL | 288.53±8.35 | 1.94 | 2.13 |
| 2 | L1 | 245.26±15.76 | 1.91 | 2.09 |
| 3 | L2 | 214.46±9.11 | 1.96 | 2.11 |
| 4 | L3 | 189.03±8.96 | 1.88 | 2.09 |
| 5 | L4 | 172.73±4.96 | 1.92 | 2.01 |
| 6 | L5 | 171.06±14.94 | 1.84 | 2.14 |
| 7 | L6 | 132.10±7.95 | 1.81 | 1.93 |
| 8 | L7 | 148.80±5.14 | 1.78 | 2.03 |
| 9 | L8 | 113.06±5.00 | 1.79 | 1.94 |

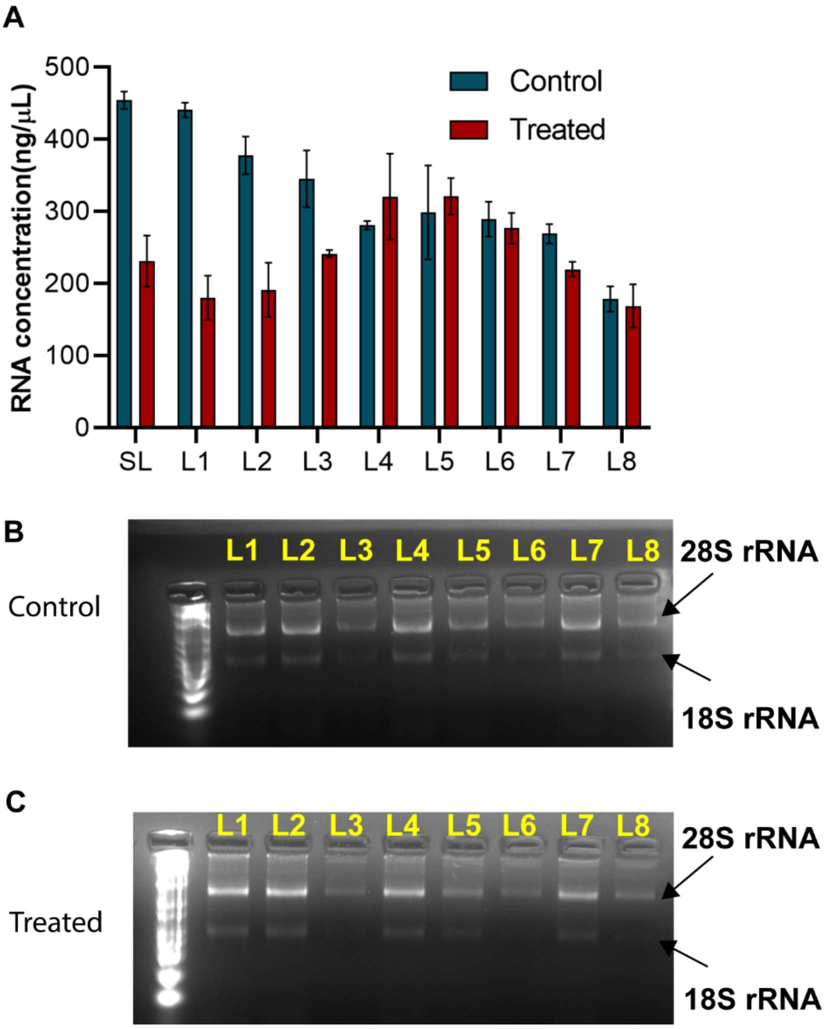


**Fig. S4. Layer-specific RNA recovery and integrity following doxorubicin treatment in multilayer paper-based 3D tumor cultures.** (A) RNA concentrations recovered from single-layer (SL) and multilayer (L1–L8) cultures under untreated (control) and doxorubicin-treated conditions. (B, C) Representative agarose gel electrophoresis images of RNA isolated from individual scaffold layers under (B) untreated and (C) doxorubicin-treated conditions, respectively. The positions of the 28S and 18S rRNA bands are indicated. RNA concentration data are presented as mean ± SD (n = 3 independent biological replicates).

**Table S7:**Purity of RNA recovered from untreated and doxorubicin-treated paper-based 3D tumor cultures across scaffold layers.

| Layer | Control A260/A280 | DOX A260/A280 | Control A260/A230 | DOX A260/A230 |
|---|---|---|---|---|
| SL | 1.96 ± 0.06 | 1.97 ± 0.01 | 2.08 ± 0.03 | 2.12 ± 0.03 |
| L1 | 1.92 ± 0.04 | 1.98 ± 0.06 | 2.07 ± 0.09 | 2.12 ± 0.04 |
| L2 | 2.05 ± 0.07 | 1.98 ± 0.12 | 1.98 ± 0.22 | 2.09 ± 0.03 |
| L3 | 1.89 ± 0.03 | 1.88 ± 0.06 | 2.03 ± 0.05 | 2.07 ± 0.04 |
| L4 | 2.09 ± 0.04 | 2.01 ± 0.04 | 2.14 ± 0.03 | 2.12 ± 0.04 |
| L5 | 1.89 ± 0.03 | 1.91 ± 0.02 | 2.13 ± 0.03 | 2.09 ± 0.03 |
| L6 | 1.90 ± 0.06 | 2.00 ± 0.05 | 2.03 ± 0.06 | 2.05 ± 0.05 |
| L7 | 2.01 ± 0.04 | 2.10 ± 0.02 | 2.17 ± 0.05 | 2.12 ± 0.03 |
| L8 | 1.98 ± 0.03 | 1.93 ± 0.07 | 2.10 ± 0.04 | 2.11 ± 0.04 |

**S1.Primer efficiency validation**

Primer efficiency validation is an essential prerequisite for reliable quantitative real-time PCR (qPCR) analysis, particularly when relative gene expression is calculated using the comparative $2^{-\Delta\Delta C_t}$ method. This method assumes that the amplification efficiencies of the target gene and the housekeeping gene are approximately equal. Significant differences in amplification efficiency can introduce systematic errors in fold-change calculations and compromise the accuracy of gene expression analysis. An amplification efficiency of 100% indicates that the PCR product doubles during each amplification cycle, while efficiencies between 90% and 110% are generally considered acceptable for quantitative analysis.

To determine primer efficiency, standard curves were generated using serial dilutions of cDNA prepared from the same RNA source used in the experimental samples. A five-point, ten-fold serial dilution series was prepared, and each dilution was amplified under identical qPCR conditions using gene-specific primers. Technical replicates and no-template controls were included to ensure reproducibility and to exclude contamination. The average Ct value obtained for each dilution was plotted against the logarithm of the initial template concentration to generate a standard curve for each primer pair (Fig. S5).

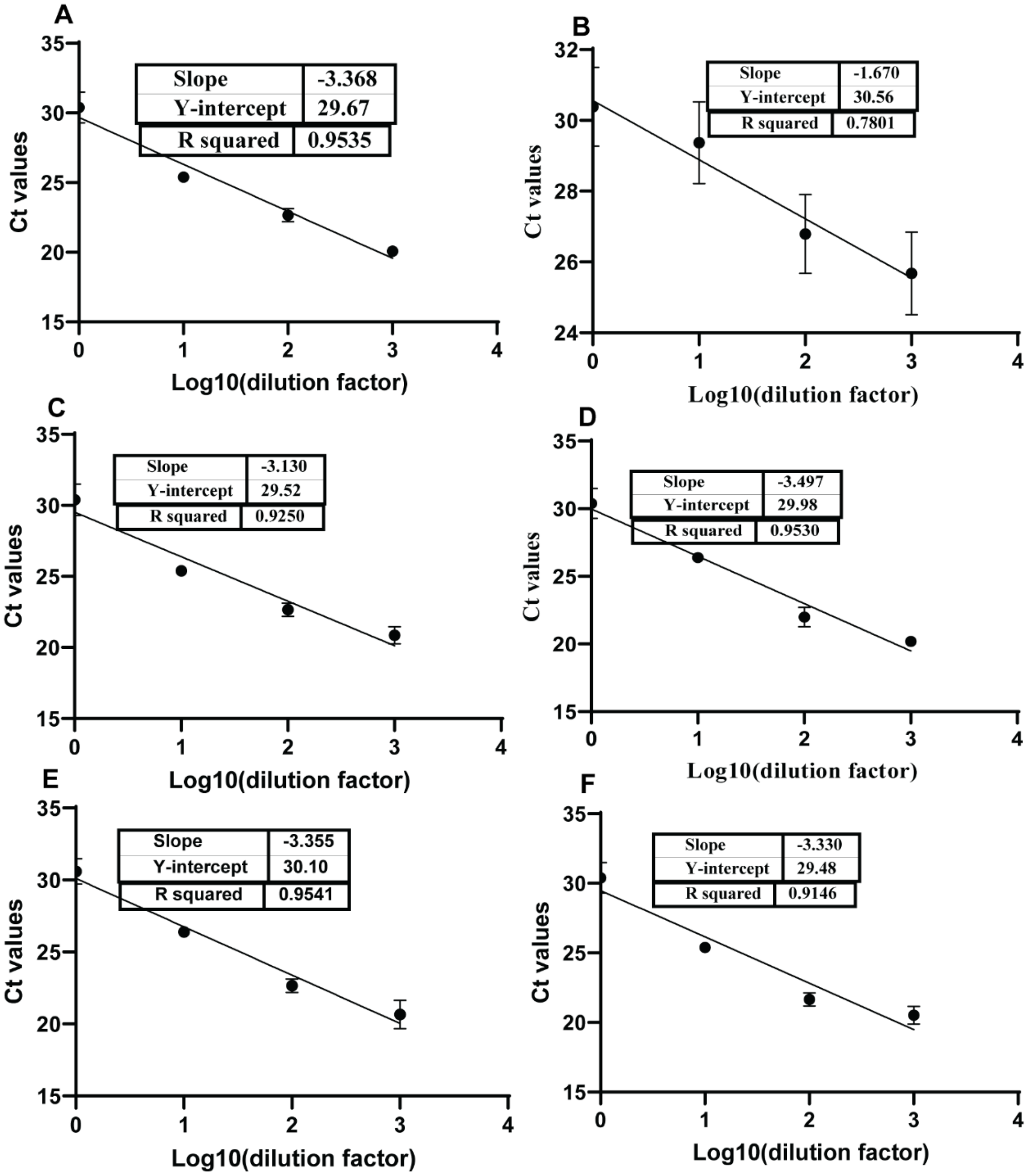


**Figure S5**: **Standard curves for primer efficiency validation.**
Standard curves obtained from serial dilutions of cDNA used to determine the amplification efficiency of the primers for (A) *GAPDH*, (B) *Actin*, (C) *GLUT4*, (D) *HIF1A*, (E) *HIF2A*, and (F) *VEGFA*. Primer efficiency was calculated from the slope of the linear regression according to Equation S1.

The amplification efficiency (**E**) for each primer pair was calculated from the slope of the regression line using Equation S1:

$$Amplification\ effeciency(\%) = (10^{(\frac{-1}{slope})} - 1) \times 100 \qquad S1$$

where the slope is obtained from the linear regression of Ct values versus the logarithm of template concentration.

The slopes obtained from the standard curves, the corresponding amplification efficiencies, and their interpretations are summarized in Table S7. The primer pairs for GAPDH, HIF-1α, HIF-2α, VEGF, and GLUT4 exhibited amplification efficiencies ranging from 93.18% to 108.68%, which fall within the generally accepted range of 90–110%, confirming their suitability for quantitative gene expression analysis using the $2^{-\Delta\Delta C_t}$ method. In contrast, the Actin primer exhibited an amplification efficiency of 297.01%, indicating non-specific amplification or primer-related artifacts. Therefore, GAPDH, which exhibited an amplification efficiency of 98.11% (Table S8), was selected as the housekeeping gene for normalization in all subsequent qPCR analyses.

**Table S8.**Amplification efficiency analysis of primers used for quantitative real-time PCR (qPCR).

| **Gene** | **Slope** | **Amplification Efficiency(%)** | **Interpretation** |
|---|---|---|---|
| Actin | –1.670 | 297.01% | Too high – likely non-specific |
| GLUT4 | –3.130 | 108.68% | Acceptable |
| VEGFA | –3.330 | 99.66% | Excellent |
| HIF2A | –3.356 | 98.60% | Excellent |
| GAPDH | –3.368 | 98.11% | Excellent |
| HIF1A | –3.497 | 93.18% | Good |

**Table S9. Dunnett's multiple comparisons test for cell viability following one-way ANOVA.**

| **Comparison** | **Mean Difference** | **95% Confidence Interval** | **Adjusted *P* value** | **Significance** |
|---|---|---|---|---|
| **L1 vs. L2** | 11.60 | −7.113 to 30.32 | 0.3474 | ns |
| **L1 vs. L3** | 18.27 | −0.4504 to 36.99 | 0.0573 | ns |
| **L1 vs. L4** | 40.54 | 21.82 to 59.25 | <0.0001 | **** |
| **L1 vs. L5** | 55.50 | 36.78 to 74.22 | <0.0001 | **** |
| **L1 vs. L6** | 54.72 | 36.00 to 73.43 | <0.0001 | **** |
| **L1 vs. L7** | 58.30 | 39.58 to 77.01 | <0.0001 | **** |
| **L1 vs. L8** | 54.29 | 35.57 to 73.01 | <0.0001 | **** |

**Results of Dunnett's multiple comparisons test following one-way ANOVA for gene expression analysis.**

**Table S10.** ***HIF1A*** **gene expression**

| Comparison | Mean Difference | 95% Confidence Interval | Adjusted *P* value | Significance |
|---|---|---|---|---|
| L1 vs. L2 | -0.3200 | -0.5785 to -0.0615 | 0.0125 | * |
| L1 vs. L3 | -0.5870 | -0.8455 to -0.3285 | <0.0001 | **** |
| L1 vs. L4 | -0.8203 | -1.0790 to -0.5618 | <0.0001 | **** |
| L1 vs. L5 | -1.1640 | -1.4230 to -0.9058 | <0.0001 | **** |
| L1 vs. L6 | -1.3320 | -1.5910 to -1.0730 | <0.0001 | **** |
| L1 vs. L7 | -1.6850 | -1.9440 to -1.4260 | <0.0001 | **** |
| L1 vs. L8 | -1.6520 | -1.9110 to -1.3930 | <0.0001 | **** |

**Table S11.** ***HIF2A*** **gene expression**

| **Comparison** | **Mean Difference** | **95% Confidence Interval** | **Adjusted *P* value** | **Significance** |
|---|---|---|---|---|
| L1 vs. L2 | -0.1200 | -0.4768 to 0.2368 | 0.8603 | ns |
| L1 vs. L3 | -0.6358 | -0.9926 to -0.2790 | 0.0005 | *** |
| L1 vs. L4 | -0.5962 | -0.9530 to -0.2394 | 0.0010 | *** |
| L1 vs. L5 | -1.2540 | -1.6110 to -0.8975 | <0.0001 | **** |
| L1 vs. L6 | -1.4020 | -1.7590 to -1.0450 | <0.0001 | **** |
| L1 vs. L7 | -1.7860 | -2.1430 to -1.4290 | <0.0001 | **** |
| L1 vs. L8 | -1.9880 | -2.3440 to -1.6310 | <0.0001 | **** |

**Table S12.** ***VEGFA*** **gene expression**

| Comparison | Mean Difference | 95% Confidence Interval | Adjusted *P* value | Significance |
|---|---|---|---|---|
| L1 vs. L2 | -0.2500 | -0.8974 to 0.3974 | 0.7756 | ns |
| L1 vs. L3 | -0.5419 | -1.1890 to 0.1055 | 0.1219 | ns |
| L1 vs. L4 | -0.9336 | -1.5810 to -0.2862 | 0.0037 | ** |
| L1 vs. L5 | -1.1420 | -1.7900 to -0.4949 | 0.0006 | *** |
| L1 vs. L6 | -1.2330 | -1.8800 to -0.5851 | 0.0003 | *** |
| L1 vs. L7 | -1.6560 | -2.3030 to -1.0080 | <0.0001 | **** |
| L1 vs. L8 | -1.7300 | -2.3770 to -1.0820 | <0.0001 | **** |

**Table S13.*GLUT4*** **gene expression**

| Comparison | Mean Difference | 95% Confidence Interval | Adjusted *P* value | Significance |
|---|---|---|---|---|
| L1 vs. L2 | -0.0123 | -0.0776 to 0.0530 | 0.9912 | ns |
| L1 vs. L3 | 0.0346 | -0.0307 to 0.1000 | 0.4990 | ns |
| L1 vs. L4 | 0.0644 | -0.0009 to 0.1297 | 0.0542 | ns |

| | | | | |
|---|---|---|---|---|
| L1 vs. L5 | 0.1310 | 0.0657 to 0.1963 | 0.0002 | *** |
| L1 vs. L6 | 0.2440 | 0.1787 to 0.3093 | <0.0001 | **** |
| L1 vs. L7 | 0.1977 | 0.1324 to 0.2630 | <0.0001 | **** |
| L1 vs. L8 | 0.1904 | 0.1251 to 0.2557 | <0.0001 | **** |

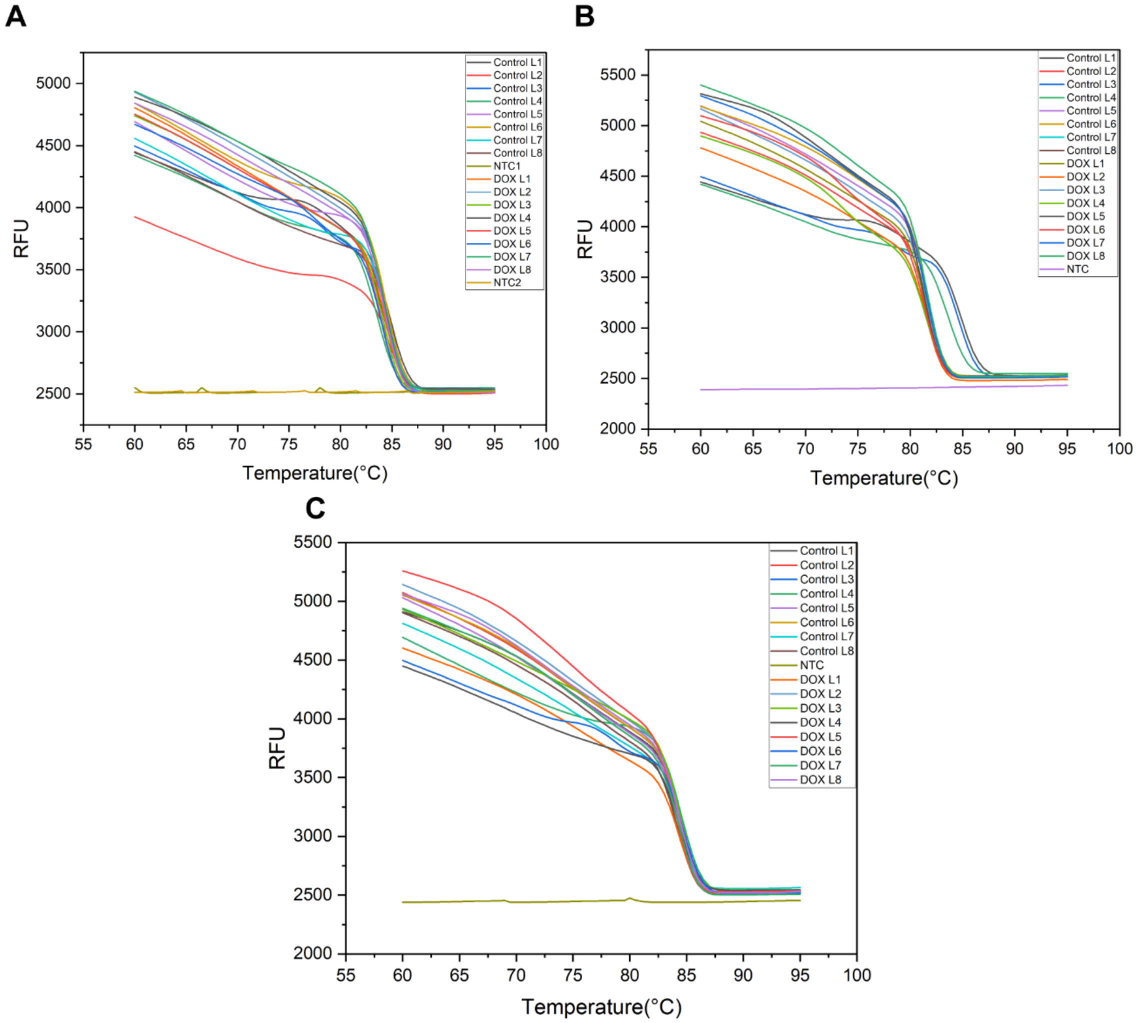


**Fig. S6. Melting-curve analysis of qRT-PCR products from untreated and doxorubicin-treated multilayer paper-based 3D tumor cultures. Fluorescence melting profiles (RFU versus temperature) for** (A) *GAPDH*, (B) *HIF1A*, and (C) *VEGFA* across scaffold layers L1–L8 under untreated (control) and doxorubicin-treated conditions. No-template controls (NTCs) are included to assess background amplification.

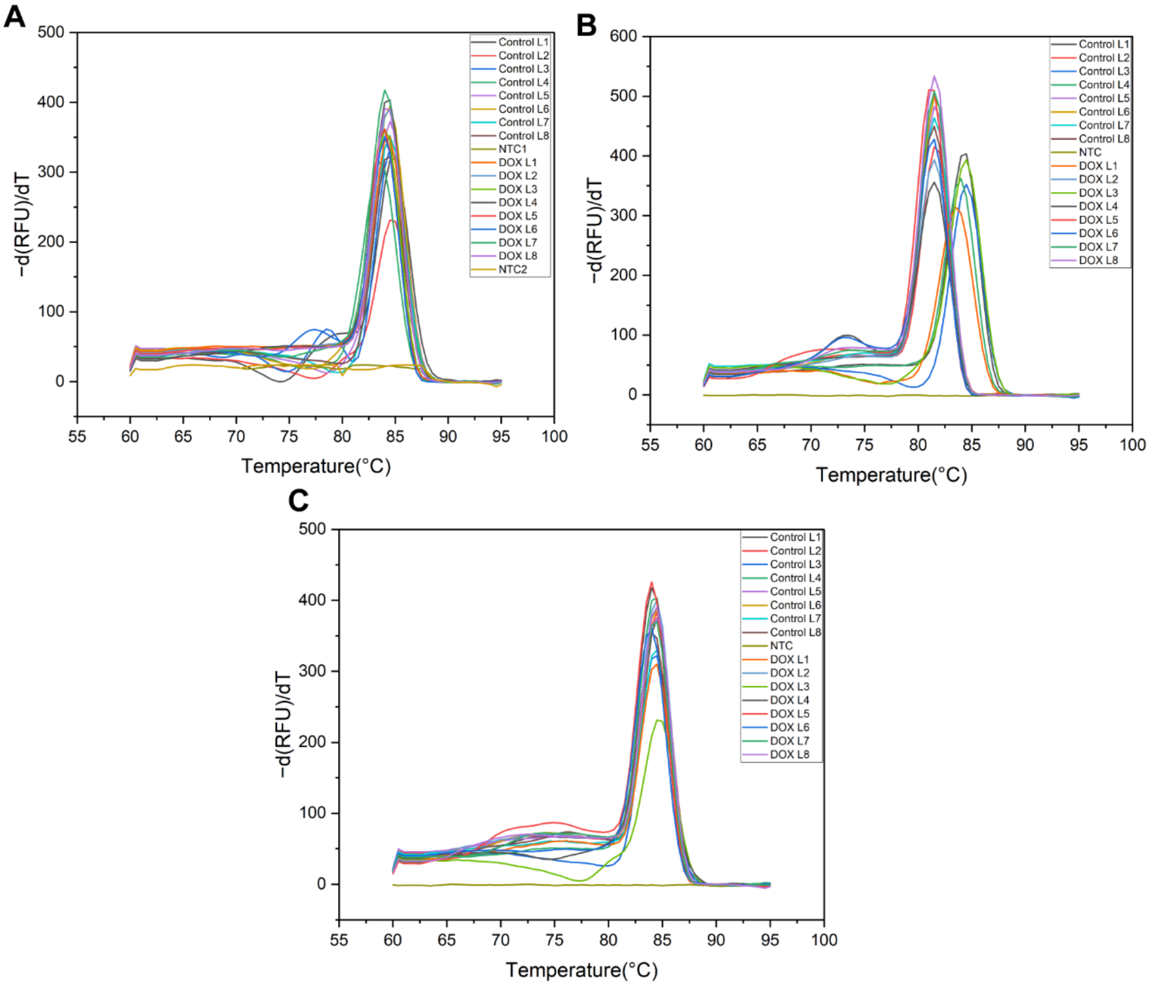


**Fig. S7. Derivative melt-curve analysis of qRT-PCR products from untreated and doxorubicin-treated multilayer paper-based 3D tumor cultures. Derivative melting curves (−dRFU/dT) for** (A) *GAPDH*, (B) *HIF1A*, and (C) *VEGFA* across scaffold layers L1–L8 under untreated (control) and doxorubicin-treated conditions. No-template controls (NTCs) are included to assess background amplification.

**Table S14:**Statistical significance of DOX-induced changes in *HIF1A* and *VEGFA* expression across scaffold layers relative to the no-change reference value.

| **Layer** | **HIF-1α p-value** | **Significance** | **VEGFA p-value** | **Significance** |
|---|---|---|---|---|
| **L1** | 0.055205 | ns | 0.035601 | * |
| **L2** | 0.037956 | * | 0.036653 | * |
| **L3** | 0.169966 | ns | 0.083763 | ns |
| **L4** | 0.086193 | ns | 0.056023 | ns |
| **L5** | 0.004913 | ** | 0.041870 | * |
| **L6** | 0.029535 | * | 0.033923 | * |
| **L7** | 0.036947 | * | 0.020027 | * |
| **L8** | 0.005229 | ** | 0.000570 | *** |
